# Approximate local alignment via chained MEM divergence estimation for detecting horizontal gene transfer

**DOI:** 10.64898/2026.10.01.756044

**Authors:** Grace Oualline, Sakshi Pandey, Xiaolei Brian Zhang, Christina Boucher, Yun William Yu

**Affiliations:** Carnegie Mellon University; University of Florida; New York Genome Center

## Abstract

Local alignment tools struggle at querying against thousands of bacterial genomes, because the reference indexes are memory-intensive and base-level dynamic programming is slow. We present alamem (https://github.com/yunwilliamyu/alamem), an approximate local aligner that streams unprocessed reference databases and estimates average nucleotide identity (ANI) from Maximal-Exact-Match (MEM) chaining statistics. Alamem can query a bacterial genome against >85,000 prokaryotic genomes in under 30 seconds (with multithreading) using 4 GB of memory, making it over 16x faster and 21x more memory-efficient than existing local aligners using their default running modes. On top of alamem, we built alasight (https://github.com/graceoualline/alasight) for detecting signatures of horizontal gene transfer (HGT) by finding pairs of alamem hits to divergent genomes. Alasight achieves over an order of magnitude better sensitivity on data from real bovid microbiomes than existing mobile genetic element-detection software and provides depth and sparsity information supporting each putative HGT region.

## Main Text

One of the fundamental operations in bioinformatics is local sequence alignment, which is widely used to identify regions of sequence similarity that may indicate biological homology—that is, whether they share a common ancestor. This problem was addressed in 1981 with the development of Smith-Waterman dynamic programming^1^, and has been succeeded by many approximate heuristics over the decades, including NCBI BLAST^2^, BLAT^3^, LAST^4^, FastGA^5^, and minimap2^6^. However, applying existing local alignment methods to modern datasets spanning hundreds of thousands of genomes is computationally prohibitive, and there remains a need for scalable frameworks capable of operating at this magnitude.

A motivating application demanding this scale is the detection of mobile genetic elements (MGEs) in prokaryotic genomes. MGEs are genetic elements capable of moving within or between genomes. Their intercellular movement drives horizontal gene transfer (HGT) across bacterial species. This trafficking accelerates bacterial evolution and is a principal route by which novel antibiotic resistance genes (ARGs) spread through microbial communities^7–8^. Current MGE detection tools are constrained by their narrow focus on specific MGE classes and a reliance on curated databases. While specialized pipelines exist for MGEs such as integrative and conjugative elements (ICEs), transposons, insertion sequences (ISs), plasmids, and integrons, these approaches depend on signature protein databases^9^, or curated profiles used with hidden Markov models to detect specific modules^10^. This reliance is a common theme across several state-of-the-art tools. MobileElementFinder aligns queries to a database of known MGEs^8^. geNomad classifies sequences as plasmid or virus using a model trained on annotated genomic databases^11^. As bacterial genomes evolve rapidly, and ARGs continue to emerge, the ability to detect MGEs without dependence on a reference database becomes increasingly critical.

An early reference-free approach for detecting novel HGT was pioneered in Alien Hunter^12^, which flags regions of DNA compositionally different from their surroundings. While useful, compositional signatures can fade over time; a more reliable approach is to leverage evolutionary history. A high-similarity sequence shared between two species that diverged millions of years ago, and that is not found in other related species, likely did not arrive there by descent, but by jumping between the species via HGT. We previously introduced skandiver, which finds genomic regions that align to divergently different species^13^. Skandiver finds distinctive evolutionary signatures using skani^14^ for whole-fragment ANI computation. But, skani computes *global* similarity of candidate sequences. For local alignment, sequences must be first partitioned, preventing skandiver from being able to resolve the precise boundaries of the elements it detects. Unfortunately, we found existing local aligners to be either too slow or memory-intensive to search over the entire universe of bacterial genomes.

This led us to build alamem (Approximate Local Alignment using MEMs), overcoming two key algorithmic and engineering problems along the way. First, existing local aligners have an expensive base-level dynamic programming (DP) extension step to resolve boundaries and similarity. Skani avoids extension via k-mer subsampling for global similarity, but local alignment requires precise, base-level comparison that subsampling cannot recover. Here, we resolve this problem by combining insights from probabilistic sketching^14^ and matching statistics^15^. More precisely, by chaining MEMs^16–17^ instead of k-mers, we obtain more precise boundary resolution in the high similarity regime without incurring additional computational overhead. Then, using an average-case analysis on a point-mutation model^18^, we estimate ANI from the average MEM length, the aligned gap length, and the unaligned gap length, which avoids DP extension.

Second, existing local aligners are designed around building an index of the reference genome(s) in memory. For the Genome Taxonomy Database^19^ r214 representative genome set, which is around 550 gigabases of sequence across 85,205 genomes, this would require hundreds of gigabytes to terabytes of RAM, even with modern compressed data structures. Furthermore, even with a prebuilt index, we still need time to load the index into RAM, which is an amount of data proportional to the full size of the database. To reduce memory requirements, we reverse the usual design by indexing the query and streaming the (unindexed) reference sequences instead. In our regime, the query (e.g., a single bacterial genome) is small enough to easily fit into RAM, so we use a conventional flat hash table for speed, avoiding cache inefficiencies that have been traditionally associated with compact data structures.

Alamem is designed for a use case similar to BLAT’s, finding short matches (>40 bp) of >90% ANI across entire bacterial genomes, though the default parameters are tuned for >96% ANI. On a 100 kbp query against a small dataset of 1,000 prokaryotic genomes, alamem is ~16x faster than BLAT in single-threaded mode, achieving performance superior even to modern aligners such as minimap2. Furthermore, the memory requirements for this experiment are a modest 200 MB (Figure 1). Using 64 threads, querying a 2 Mbp bacterial genome against the full 85,205-genome GTDB r214 representatives dataset takes <30 seconds and 4 GB of RAM.

**Figure 1:**
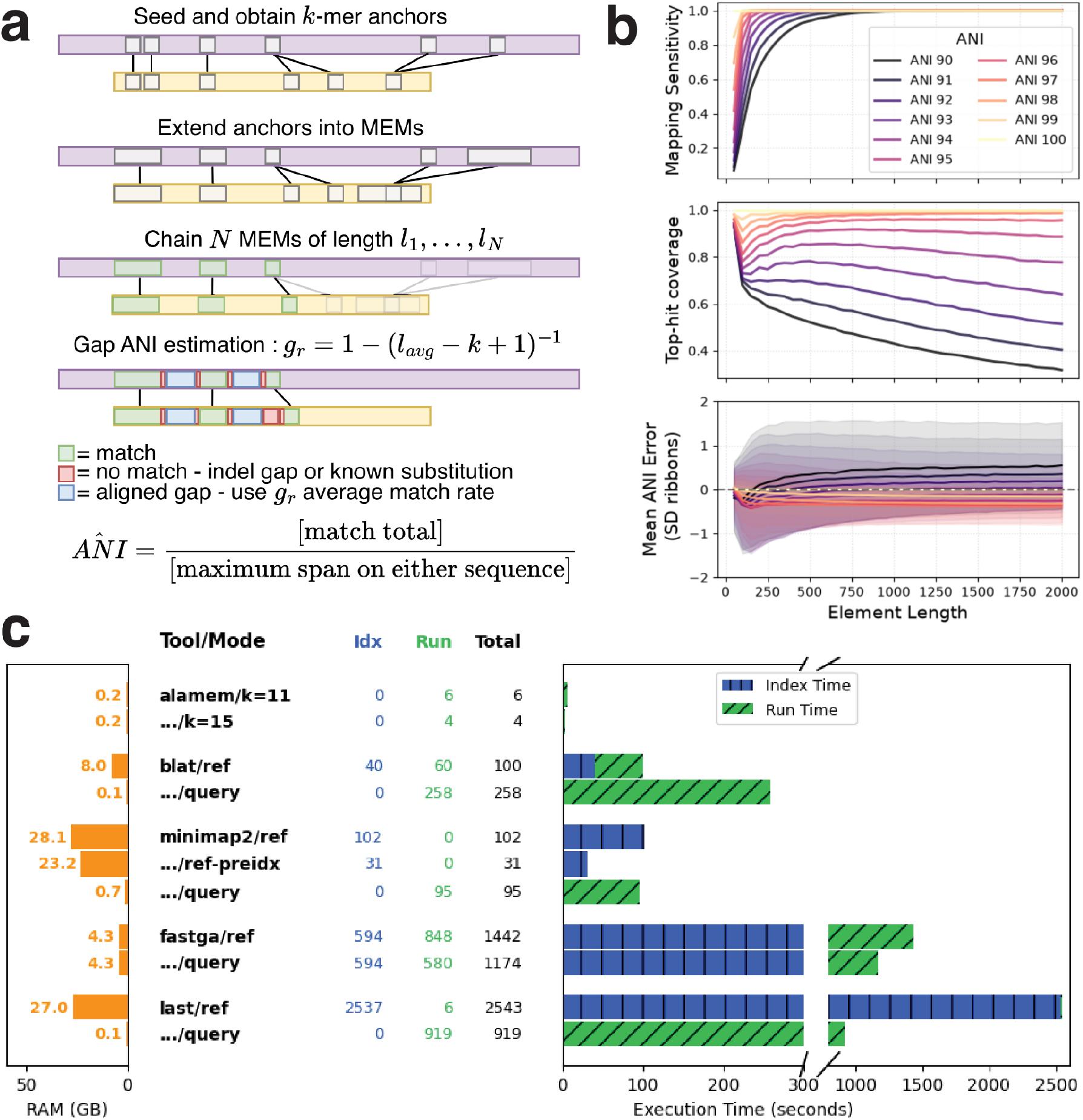
**(a)** alamem works by seeding k-mer anchors, exact extension into Maximal-Exact-Matches (MEMs), chaining those MEMs, and then directly estimating ANI from MEM length and gap lengths. **(b)** On simulated fragments of length 50 to 2000 bp drawn from a 100 kbp reference: (top) alamem with k=15 and min_ani=90% recovers nearly all fragments; (middle) hits cover most of each fragment, except when the true ANI is near the min_ani threshold; (bottom) mean ANI error is typically <1% (see Supplementary Figures 1-4 for k=11/13 and Pearson correlation scatterplots). **(c)** alamem is faster and uses less memory; numbers from querying a 100 kbp fragment against 1,000 bacterial genomes (total sequence length just under 5 Gbp) taken from the GTDB (<1% of the GTDB representative genomes by sequence). Because alamem indexes the query rather than the reference database, competing methods were additionally evaluated with the query indexed instead of the reference, which typically lowered memory requirements at the cost of runtime (see Supplementary Figure 5 for alamem’s near-linear scaling with respect to database size and thread count).

Leveraging alamem, we developed alasight (ALAmem for SIGnatures of Horizontal Transfer), a scalable HGT discovery method capable of uncovering novel elements absent from MGE catalogs. To detect transfer events, alasight applies an overlap-divergence filter requiring that a genomic region align to at least two divergent species, which are themselves divergent from the query sequence. Since GTDB r214 is already deduplicated, this more precisely amounts to requiring two different hits that are ≤95% ANI to the query sequence. Further, alasight displays the depth of support (number of divergent clades that support a hit) and sparsity of support (fraction of leaves within the smallest subtree of the TimeTree of Life^20^ containing them). These metrics distinguish vertically conserved elements from true HGT, which is expected to result in multiple sparsely spread out hits across the TimeTree of Life (Figure 2).

**Figure 2:**
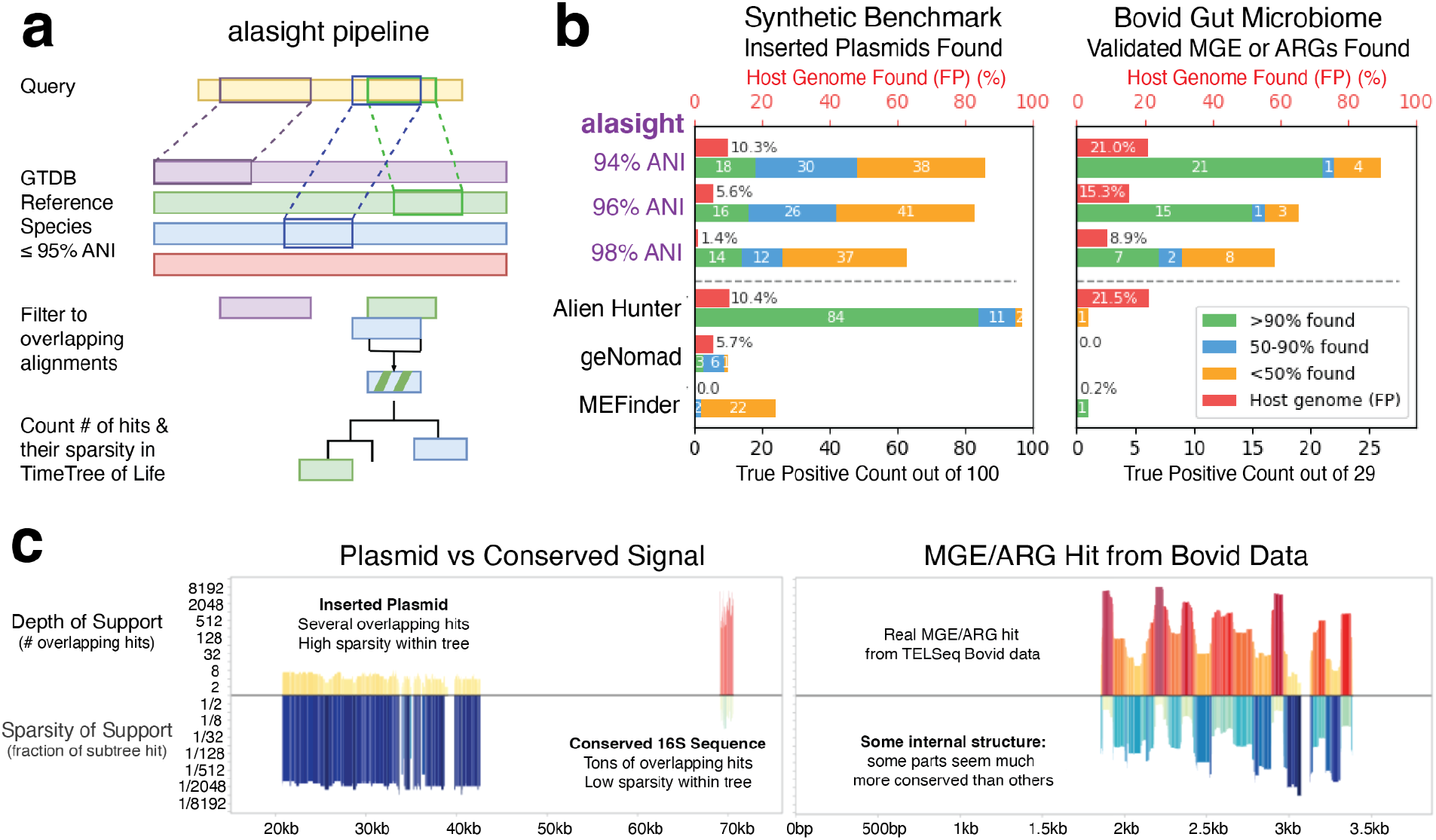
**(a)** Alasight uses alamem (with k=15) to find overlapping hits to divergent GTDB reference genomes (≤95% ANI). A minimum depth of two hits is required, but alasight presents depth of support (# hits) and sparsity of support (fraction of subtree in TimeTree of Life with hits) as signal. **(b)** On (left) simulated benchmarks using plasmids inserted into prokaryotic genomes, and (right) 13 real TELSeq consensus sequences from a bovid gut microbiome with TELCoMB independently validated MGEs and ARGs (total of 29), alasight performs well compared to Alien Hunter, geNomad, and MobileElementFinder (MEFinder) at recovering hits; only Alien Hunter with the inserted plasmids did better, but it was unable to be run on the short TELSeq consensus sequences for all but one sequence. On that one (long) sequence, Alien Hunter recovered <50% of the annotated MGE/ARG but had such a high FP rate (46%) that the overall FP% across all host genomes was 21.5% (see Supplementary Figure 6). **(c)** (left) Simulated distinction between conserved elements vs inserted plasmid, showcasing the different signatures of depth and sparsity. (right) Real alasight result for one of the TELSeq MGE/ARG hits, showcasing potential internal structure. For depth, we record the number of clade hits that support each region of the genome. For sparsity, we find the smallest subtree of the TimeTree of Life that contains all hits, and measure the fraction of hit leaves over total leaves in that subtree.

On a synthetic benchmark we built of plasmids inserted into bacterial hosts, alasight’s recovery to false positive (FP) rates are only slightly worse than Alien Hunter in the high FP regime and better than geNomad and MobileElementFinder in the medium and low FP regimes respectively. On real consensus sequences from a cow gut resistome generated by target-enriched long-read sequencing^21–22^ with 29 annotated MGE or ARG regions, alasight recovered between 17 and 26, depending on FP regime, while none of the other tools were able to recover more than 1 (Figure 2). We further note that some of those FP hits may well be real instances of HGT; MGEs frequently co-transfer adjacent chromosomal and accessory genes that fall outside strict reference annotations^7^. Providing such candidates for downstream discovery is an important application of alasight, which we hope will help biologists shed light on such phenomena.

In conclusion, alamem is over an order of magnitude faster than existing local alignment heuristics, while recovering over 90% of high-ANI hits. It does this by combining insights from probabilistic sketching and matching statistics to avoid base-level extension for similarity computation. Using alamem, our alasight software is able to detect signatures of horizontal gene transfer without using reference MGE databases, outperforming existing methods, especially on short sequences.

## Online Methods

### Alamem methodological details

Alamem starts by indexing all of the canonical k-mers within the query using a flat hash table, marking their locations. We assume that the query fits in memory, which is reasonable for bacterial genomes, contigs, and consensus sequences on the order of 1–100 Mbp. Optionally, frequent k-mers can be masked, but this is off by default. Alamem also puts a 4 MiB 1-bit Bloom prefilter in front of the index to guard against cache misses from missing k-mers.

When alamem streams the reference genomes across the index, it defaults to the same strategy used in BLAT, whereby we use equal k-mer length and stride. If a k-mer is invalid (due to the presence of an N), we deliberately skip the entire k-mer, rather than starting over after the N in the sequence. This strategy allows for easy vectorization by the compiler and better pipelining, as there is very little branching in the code.

Canonical k-mer matches between the reference and query form the set of initial k-mer anchors. We then prune any isolated anchors that cannot be extended or chained into a hit of a given minimum length. The remaining anchors are then extended maximally to both the left and the right as Maximal-Exact-Matches (MEMs), avoiding any anchors that lie entirely within an already extended MEM, as their extensions would be identical.

Next, we prune isolated short MEMs that cannot be extended or chained into a hit of minimum length (like we did with anchors beforehand). Multiple pruning steps are used to reduce the candidates passed to the computationally expensive sorting and chaining stage. All the MEMs are sorted by the sum of their start points on the query and reference (anti-diagonal sorting), and for chaining, we have a lookback window bounded by twice the maximum gap allowed (with a hard-coded heuristic lookback bound of 5,000 MEMs to deal with super repetitive regions and long MEMs). Chaining only happens for collinear MEMs with gaps below the maximum gap allowed; the anti-diagonal sort order ensures that collinear predecessors are always in the lookback.

The scoring function we use for chaining follows the form of the final ANI computation below, but instead uses the minimum ANI allowed for hits so that it is a simple scoring function. The chaining score adds the number of matches from the MEM itself plus the number of expected matches from [(aligned_gap_size-2) x minimum_ANI] and subtracts any unaligned gap.

Unaligned gap size is the absolute difference in gap sizes between the gaps on the reference and the query, which can only be a result of indels, and so cannot contribute to ANI. On the other hand, aligned gaps are everything else, or equivalently the minimum of the two gap sizes on the reference and query. For these, we assume that—other than the endpoints, which are known mismatches—the rest of the gap has an ANI of our minimum ANI.

After chaining, we then do a more careful evaluation of the chain ANI. Similar to above, we estimate total matches by taking all the MEM matches, and then multiplying the aligned gap length (minus the endpoints) by a regional ANI. The regional ANI is estimated using geometric run-length theory by taking the average length of all the MEMs in the chain and subtracting out the k-mer length; this adjusted average MEM length can be transformed into an estimator for ANI because the lengths are exactly the lengths of runs of successes in a sequence of Bernoulli trials with parameter regional ANI. The adjustment is necessary because we have to condition out finding the MEM to begin with, which we do using exact k-mer matching, so the first k successes are guaranteed. The denominator for the final ANI calculation is the total aligned span, which penalizes properly for any indels. This final ANI is not an unbiased estimator because the MEM and gap lengths are not actually randomly sampled, but is pretty close otherwise.

More precisely, imagine we are in a regime with mutation rate θ and ANI Δ = 1 − θ. Then the expected length of a MEM is simply a Bernoulli run length (not counting the mismatch that breaks the run), so 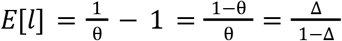. If we averaged over all MEM lengths to get *l*_*avg*_ ≈ *E*[*l*], we’d then be able to estimate *l*_*avg*_(1 − Δ) = Δ, so *l*_*avg*_ = Δ(1 + *l*_*avg*_), implying that 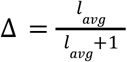. This formula however only works if *l*_*avg*_ is the average length over *all* MEMs in the chain. However, we extend *k*-mer matches to MEMs, so the only MEMs we can see are ones of at least length *k*. To correct for that, we have to shift by *k*, leading us to the formula 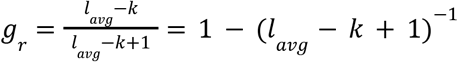, which we use as our regional gap ANI estimate in Figure 1.

There are however two problems with directly using this estimator *g*_*r*_ ≈ Δ. (1) We are assuming that all mutations, including substitutions, insertions, and deletions, are i.i.d. point mutations. This is an incorrect assumption in biological settings, as we often expect to see runs of especially insertions and deletions. (2) We also are not guaranteed to see all MEMs of length at least *k*, because we use the BLAT seeding strategy by default of looking at disjoint *k*-mers. The second problem shows up as error in our ANI estimation, which we don’t correct for because it is relatively minor when *k* is small compared to the MEM lengths. The first problem we correct by only using *g*_*r*_ as an estimator for the number of matches in aligned gaps, not counting the boundary bases, which we know do not match. Then, we simply sum together the total matches (real matches from MEMs and imputed matches from aligned gaps), and divide it by the maximum span on either sequence to get our final ANI estimation.

### alamem speed and RAM benchmarking details

To compare speed and RAM requirements across methods, we took 1,000 bacterial genomes from the GTDB r214 representative genomes (out of a total of 85,205 genomes) and concatenated them into an uncompressed FASTA file. Of note, most modern tools, including alamem, can run directly on gzipped FASTA files, but that just introduces decompression time jitter. Furthermore, alamem when used with alasight prefers a text file list of genome FASTAs, rather than concatenating them, but several tools did not support that format. The total amount of sequence was just under 5 Gbp, compared to a total of around 550 Gbp in the full set of representative genomes, so less than 1% of the total sequence. Restricting ourselves to this smaller subset was necessary because otherwise most tools would run out of memory or take prohibitively long, even on our 64-core, 500 GiB RAM server. This also allowed us to run in single-thread mode, giving an apples-to-apples comparison. Of note, older tools like BLAT do not parallelize well, so the single-threaded comparison is somewhat biased in their favor.

For the query, we used a 100 kbp substring of NZ_CP013494.1, from positions 1228656-1328655. This corresponds to the plasmid pRspN6212d from *Rhizobium* sp. N6212, a member of a genus of nitrogen-fixating bacteria.

For each method (besides alamem), we ran each in both normal and “reverse” mode, where in normal mode, the tool indexed the reference genomes. In “reverse” mode, we had the tool index the 100 kbp query instead. Alamem is designed specifically to work in “reverse” mode, where we expect the query to be much smaller than the reference for speed purposes. Thus, alamem is designed such that the hits output are largely agnostic to which sequence is marked as the query or reference. Existing tools are not designed to work in reverse mode, as they make assumptions about what constitutes a good hit based on properties (such as length, or k-mer multiplicity) of the reference, so we also benchmarked them in reverse mode for completeness, which in many cases used much less RAM at the cost of runtime.

For every method, we also ran them twice in a row so that all of the reference genomes and any prebuilt indexes would be stored in the Linux RAM cache, and took the minimum of the two times, ensuring that we were not just measuring the disk read bottleneck. On our benchmark server, the working drives are spinning hard drives, and in multithreaded mode, alamem and several other tools quickly became I/O bottlenecked. These speeds are thus a proxy for the speeds when the reference is stored on fast media (such as an M.2 NVMe SSD).

For alamem, we tested with both k=11 and k=15. We tested at k=11 for maximum comparability with BLAT default options, and k=15 because that is what we found empirically to work best for alasight. k=15 is a little closer to what most other modern tools use, the maximum that the alamem software implementation currently supports, and also the default setting.

For BLAT, we used version 39×1. For overall time, we used blatHuge directly on the FASTA files. We also separately preindexed to gfidx and then used gfServerHuge to separate out the indexing and running time. We were fortunate that those times added up to our separate direct run of blatHuge. We note for completeness that using a 2bit representation slowed down BLAT by a factor of 3 when using blatHuge, so we opted for blatHuge with a plain FASTA file.

However, the 2bit representation did not slow BLAT down at all when using gfServerHuge. The results presented in Figure 1 are the best results we got using BLAT in any of our configurations. BLAT by default sets k=11.

For LAST, we ran everything using standard options. Of note, LAST uses a dynamic k-mer size, and on this dataset, we expect that k is approximately 19.

For FastGA, we used FAtoGDB + GIXmake to index the reference and the query. All of that time combined was summed to give our single FastGA indexing time; these times are the same for forward and reverse mode because FastGA requires indexing both reference and query. Then, we ran the FastGA binary itself in both directions. We used FastGA’s default k=40.

We also benchmarked minimap2, a leading high-performance mapper, to establish a benchmark for runtime and memory efficiency. To ensure that the comparison was fair, we used the options “-t 1-x asm20”. This sets the number of threads to 1 (-t 1) and parameters optimized to allow for up to 10-20% sequence divergence (-x asm20). Minimap2 sets k=19 with a minimizer window size of 10. Importantly, this choice of options (specifically, not setting -c or -a) also causes minimap2 to eschew base-level alignment, making it more comparable to alamem. We note that the 0 second runtime is not a typo; minimap2 impressively does all of the work in the indexing stage, at least according to its internal logs. Unfortunately, loading a precomputed index of 15 GB takes a substantial amount of time, which we show in the minimap2 preidx line. To be fair to minimap2, we also turned off statistics by setting “-v 2”, as those statistics take about 4 seconds to compute on this reference. We note that for applications involving many local alignments against the same reference, minimap2 remains the standard for high-performance mapping. Our advantage is when indexing and index-loading costs cannot be amortized, particularly when the reference is too large for its index to fit in memory.

### Alamem accuracy benchmarking details

The full benchmarking script for accuracy is in the scripts directory of the alamem repo. We simulated reads of varying lengths from a 100 kbp segment of a bacterial genome, with prespecified mutation rates (of the mutations, 70% substitutions, 15% insertions, and 15% deletions). Furthermore, we kept track of the mutations, so we could reconstruct the true ANI of local alignments later.

For the mapping sensitivity chart, we gave credit whenever alamem returned any hit within the read, so it is not surprising that as length increased, so did the sensitivity.

For the top-hit coverage, however, we measured the fraction of the read that was contained within the alamem hit. This was to ensure that alamem wasn’t just getting a tiny hit within a large area and calling that a success.

For the mean ANI error, we took all of the alamem hits and the alamem estimates of ANI, and compared those estimates against the true ANI, which we knew because we kept track of our simulated mutations. We displayed both the mean ANI error and also the standard deviation of the ANI error.

### Alasight methodological details

Alasight takes as input one or more genomic sequence(s) in FASTA file format, and optionally, species information for each sequence. It makes a prediction of the beginning and ending positions of all potential HGT in each input sequence. In this section, we give an overview of alasight, which can be summarized in four steps.

### Removing host species from comparison

If the query sequence shares over 95% ANI with a GTDB reference genome, we assume they are from the same species, and exclude that reference genome from HGT comparisons, as we do not consider transfers occurring within a species.

### Alignment to GTDB

Next, the query sequences are aligned to GTDB using alamem using default options in order to identify regions in GTDB that have high sequence similarity (by default >96% ANI) to the input sequences. We chose a default of 96% to give it some separation from the 95% ANI threshold we used to remove species from the comparison pool, but this value can be specified by the user, and we found 94%, 96%, and 98% to give slightly different reasonable true positive to false positive tradeoffs.

### Finding regions of overlap between divergent reference species

For each pair of alignments whose coordinates overlap on the query sequence, we want to ensure that the hits are from different species. We use by default the GTDB r214 reference genome set, where the sequences are already filtered to be distinct species, so no additional processing is needed. When using reference genomes that are not prefiltered, we compute the divergence time between the species using either the TimeTree of Life (setting a threshold of one million years divergence) or ANI below 95%. This step requires that the same region of the input sequence be independently mobile across two distinct evolutionary lineages, providing strong support against chance similarity.

### Post-processing

The regions of the input sequence that remain after the overlap-divergence filter often overlap with one another, despite aligning to different reference species. We combine these regions in two steps:

1. Merge. All regions whose coordinates overlap on the input sequence are merged into a single contiguous region.
2. Size filter. Merged regions shorter than 150 bp are discarded (user-configurable).

A single composite locus can contain modules which were acquired from phylogenetically distant donors, and now function together as one unit. Since alasight’s detections rely on alignments to divergent references, adjacent modules of one element often surface as separate, overlapping hits to different references. Combining these neighboring hits is a required step to consolidate and retrieve the entire composite HGT. A locus in which neighboring sub-regions independently match different divergent reference species is therefore evidence for, not against, horizontal acquisition; in such cases alasight reports the locus in full and includes the union of their supporting reference species.

The output is a summary file in which each row is one predicted HGT region in the input sequence, with its start and end coordinates, length, number of distinct reference species aligned to that region, and the average divergence time between the input sequence and its reference species.

#### Depth and sparsity visualization

Furthermore, for visualization, we separately compute a depth and sparsity file for each genomic region. For depth, we record the number of clade hits that support each region of the genome (with a minimum of two hits, since we filtered by overlap). For sparsity, we find the smallest subtree of the TimeTree of Life that contains all hits, and measure the fraction of hit species over total species in that subtree.

This allows disambiguation of HGT from highly conserved elements, as highly conserved elements are dense within their containing subtree, and typically have lots of hits, whereas HGT is typified by a moderate number of hits but higher sparsity. We provide plotting software to generate interactive HTML visualizations of these regions.

### Alasight benchmarking details

#### Synthetic Benchmarking

To measure accuracy against a known ground truth, we inserted plasmid fragments into bacterial sequences that contained no MGE detectable by MobileElementFinder or geNomad. We refer to these background sequences as host genomes. We chose bacterial hosts, since a plasmid inserted into a compositionally distant host (like a eukaryotic host) is easy for any method to flag and inflate perceived accuracy, while a plasmid inserted into a bacterial host is a more difficult and realistic test. For 100 synthetic sequences, a plasmid fragment was sampled from a normal distribution (mean = 10,000 bp, σ = 10,000 bp, range [1,000, 90,000] bp); the fragment was inserted at a random position within a ~100 kbp host segment, then analyzed with alasight. An inserted element was counted as detected if any reported region overlapped its true coordinates. The synthetic sequences are linked on the alasight GitHub, and available here (https://github.com/graceoualline/alasight_testdata).

#### TELSeq Benchmarking

To evaluate our approach on complex biological samples we utilized bovid fecal metagenomes from a TELSeq study by Slizovskiy et al.^21^ These libraries contained genomic DNA extracted from fresh fecal material of a periparturient Holstein-Friesian dairy cow. Due to recent treatment with an antibiotic, the sample carries a resistome shaped by real antibiotic selection, making it a biologically relevant test for recovery of ARGs and their associated MGEs. To establish reference ARG and MGE annotations we reran the TELCoMB protocol^22^ on each library using the default parameters. This workflow deduplicates the reads before aligning them to dedicated reference databases to detect ARGs and MGEs. TELCoMB retains an alignment only when it covers at least 80% of a reference ARG or 50% of a reference MGE. ARGs and MGEs occurring on the same read are then reported as colocalizations. The dataset yielded 13 consensus sequences with 29 annotated ARG and MGE features. We ran alasight on these consensus sequences using default parameters. Because TELCoMB annotates by reference alignment to a curated ARG/MGE database, and alasight does not, the TELCoMB annotations are an independent ground truth, and a region alasight recovers here is also found by a method that shares none of its machinery. In Supplementary Figure 6, we visualize the overlap of TELCoMB ground truth with alasight, MobileElementFinder, and Alien Hunter.

## Contributions

C.B. and Y.W.Y. supervised all aspects of the project. The alasight method was jointly conceived by G.O. and X.B.Z. under Y.W.Y.’s supervision. Alasight methods development was led by G.O., with S.P. assisting on parameter tuning, and X.B.Z. helping with compute benchmarking. S.P. determined the biological validation benchmarks under C.B.’s supervision. The alamem method and software were conceived and designed by Y.W.Y. All authors contributed to the writing of the manuscript.

## Acknowledgments

We would like to thank members of the Yu and Boucher labs for fruitful discussions. Additionally, we would like to thank Noelle Noyes for lending biological expertise.

## Data/Code Availability

Alamem is available at https://github.com/yunwilliamyu/alamem. Alasight is available at https://github.com/graceoualline/alasight. Data for reproducing alasight figures is available at https://github.com/graceoualline/alasight_testdata.

## Supplementary Material

**Supplementary Figure 1:**
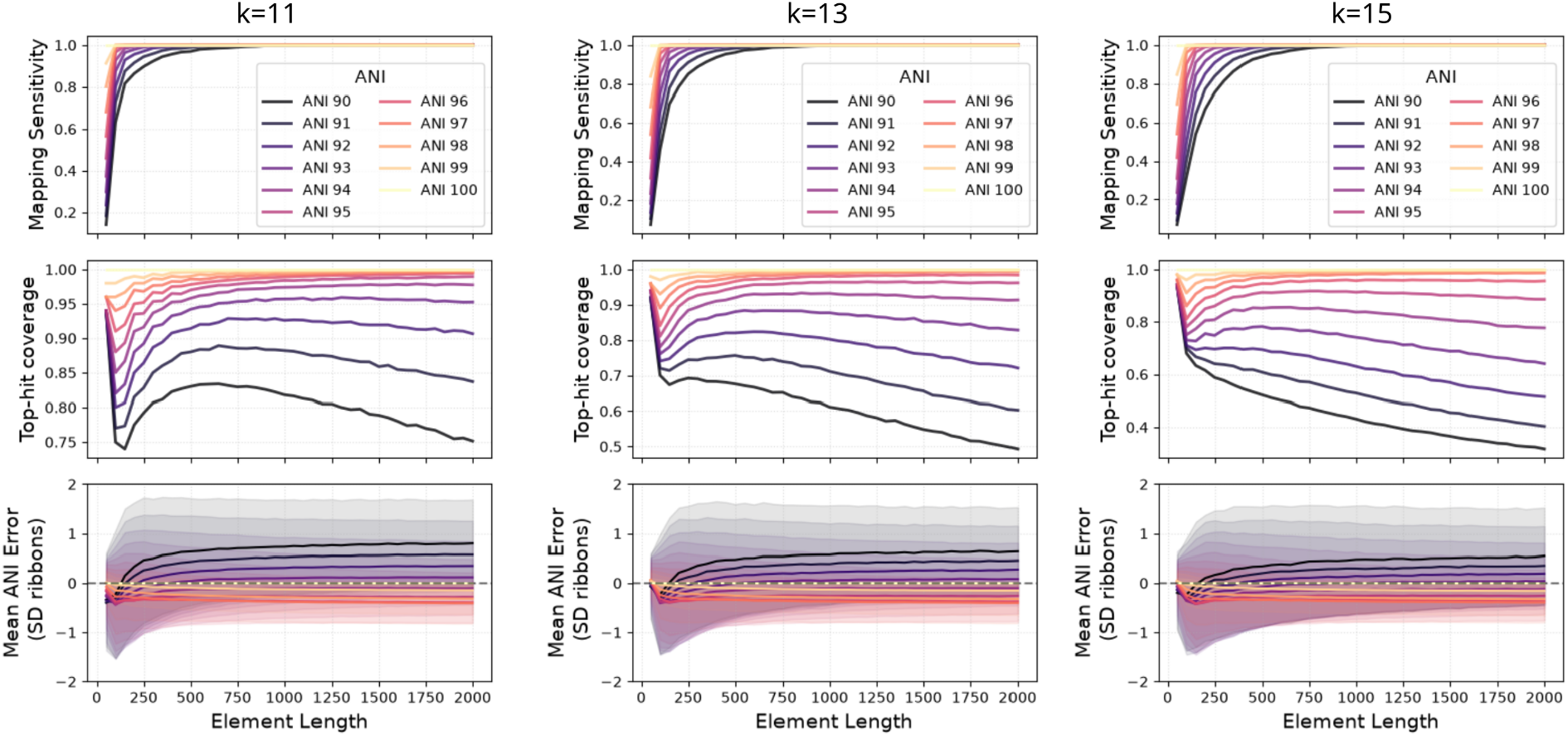
alamem’s accuracy for k=11, 13, and 15 (k=15 is a duplicate of Figure 1b). alamem’s ANI accuracy is still high even for k=13 and k=15 (the default k-value), but recovery of full low-ANI alignments suffers, and the fragments often get split into multiple hits. This largely does not matter for alasight, which operates just as well on split hits as single, unbroken ones, but setting the min_ani parameter lower can increase the contiguity of hits. Although alamem’s default min_ani is 96%, for all experiments here, we set min_ani=90%, so that we are able to accurately see accuracy for hits below 96% ANI. Notice that accuracy remains high, but the top-hit length decreases near the min_ani threshold.

**Supplementary Figure 2:**
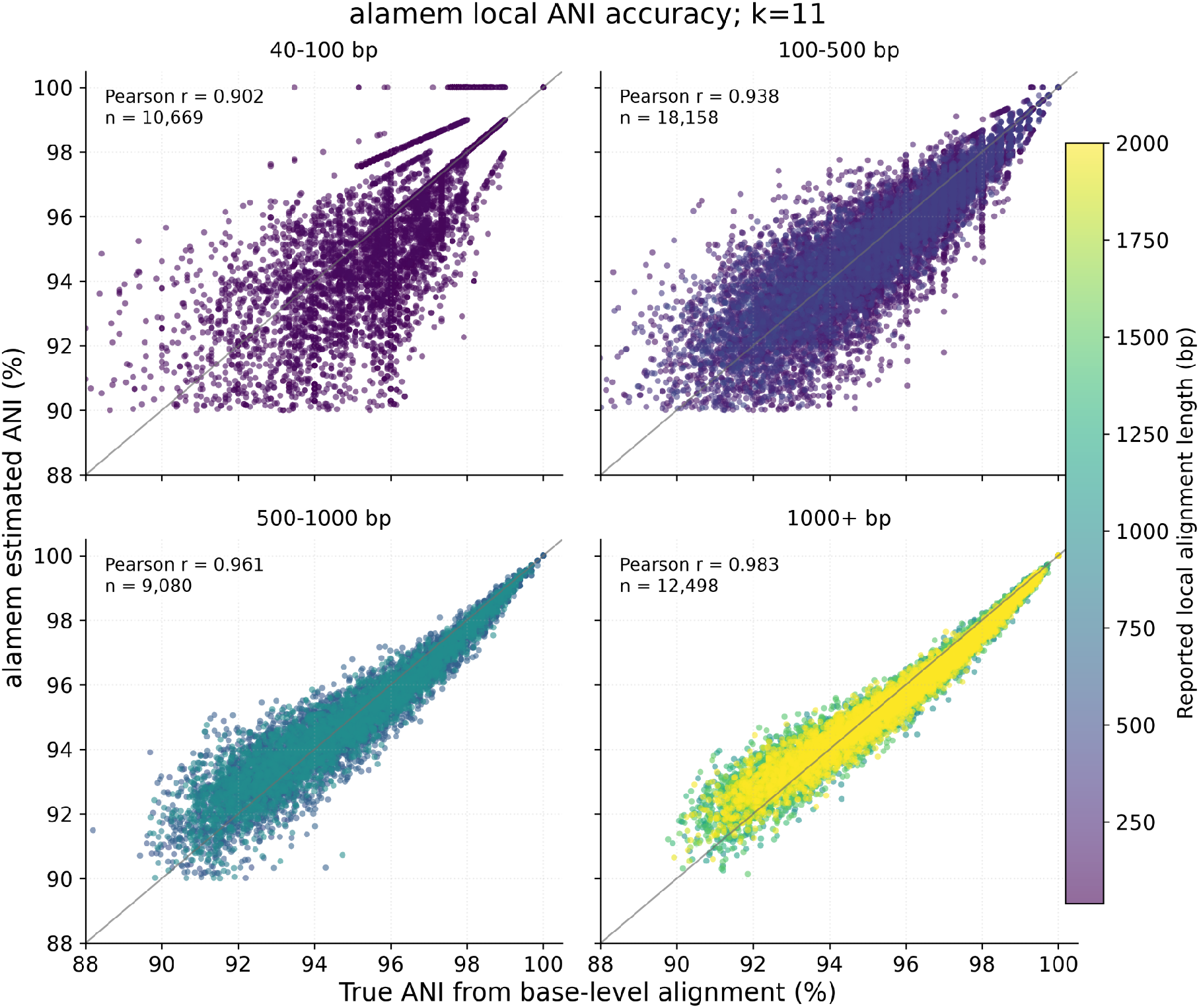
Alamem predicts ANI of local alignments with high accuracy with k=11. Comparison of true ANI to ANI estimated by alamem for individual simulated local alignments. Color indicates the reported alignment length in base pairs.

**Supplementary Figure 3:**
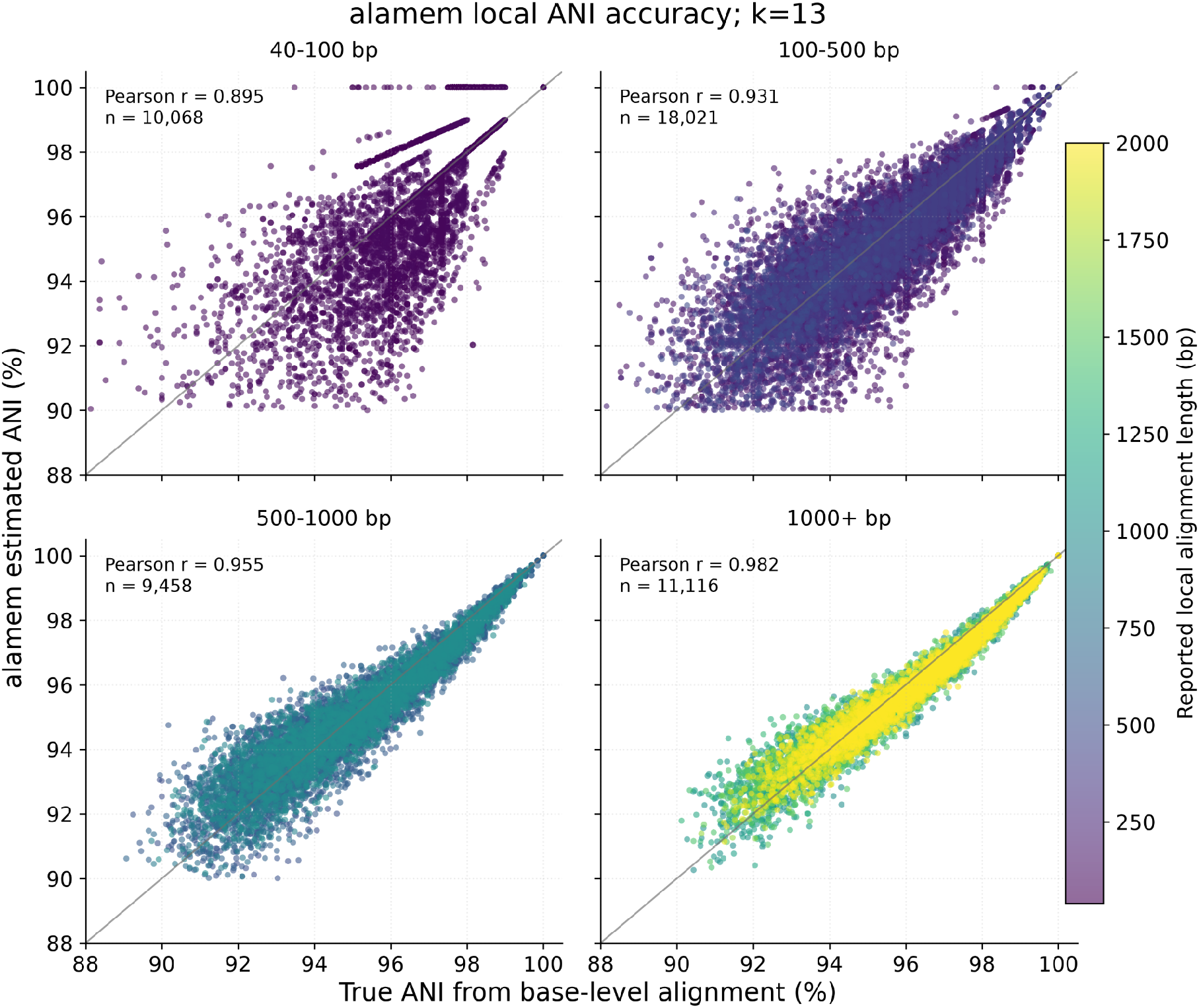
Alamem predicts ANI of local alignments with high accuracy with k=13. Comparison of true ANI to ANI estimated by alamem for individual simulated local alignments. Color indicates the reported alignment length in base pairs.

**Supplementary Figure 4:**
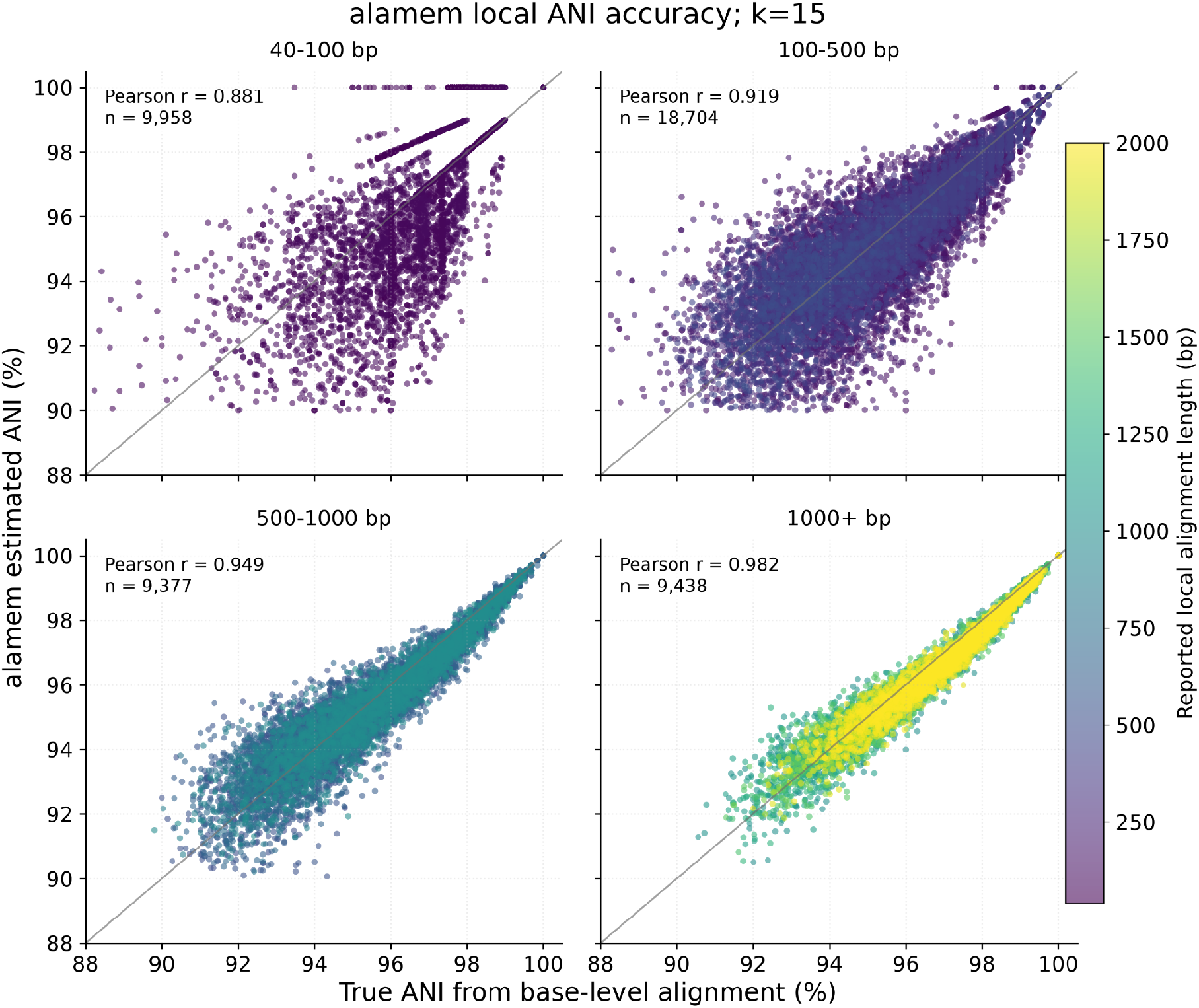
Alamem predicts ANI of local alignments with high accuracy with k=15. Comparison of true ANI to ANI estimated by alamem for individual simulated local alignments. Color indicates the reported alignment length in base pairs.

**Supplementary Figure 5:**
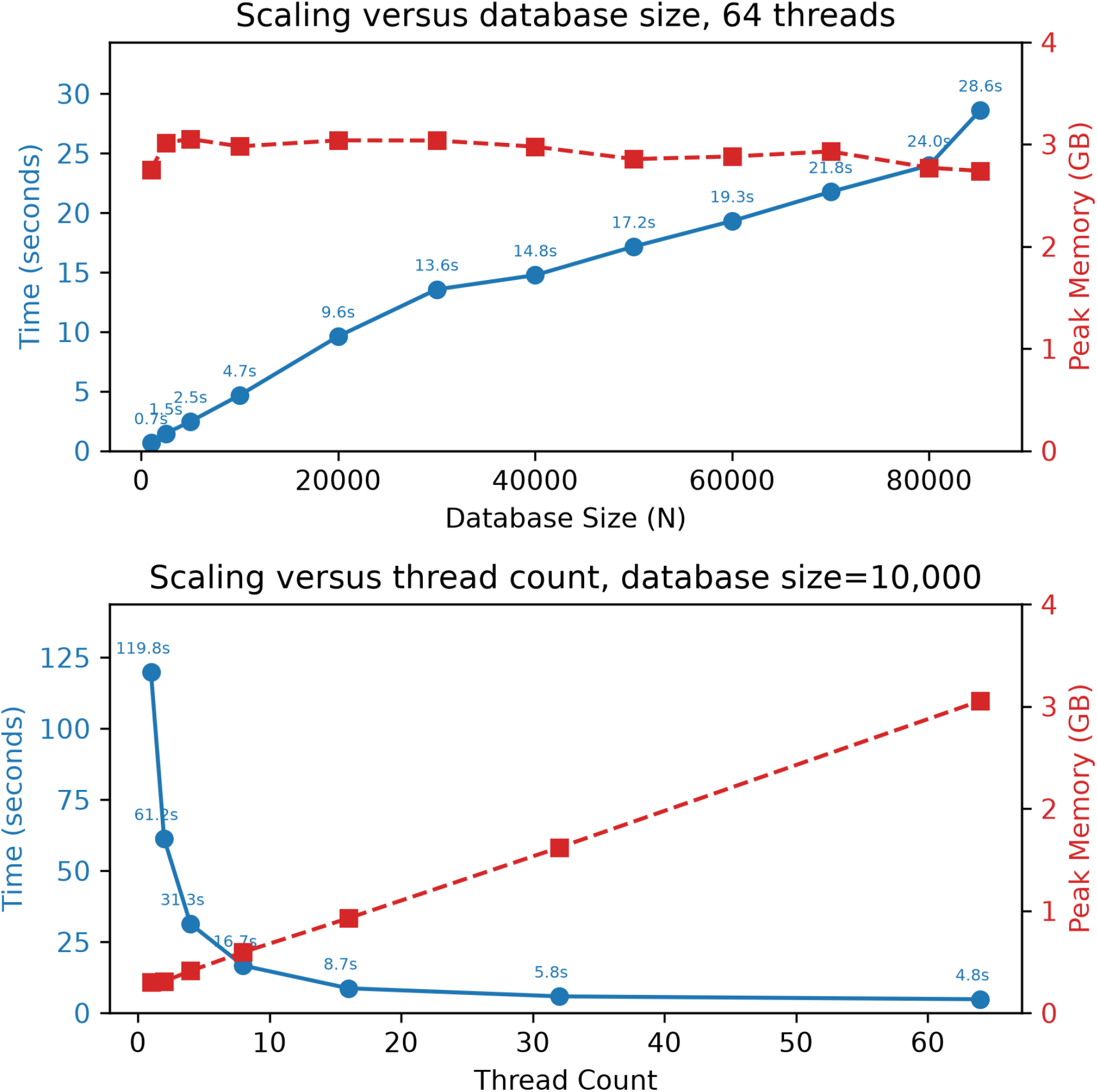
Alamem scaling performance querying a 1 Mbp subsequence of *Fusobacterium nucleatum* against GTDB r214 representative genomes. **(top)** Wall-clock runtime and peak memory usage as the searched database size increases from 1,000 to 85,205 representative genomes using 64 threads. **(bottom)** Wall-clock runtime for searching 10,000 representative genomes as the thread count increases from 1 to 64. Benchmarks were run on a dual-socket Intel Xeon Gold 5416S system with 64 physical cores and 500 GiB RAM.

**Supplementary Figure 6:**
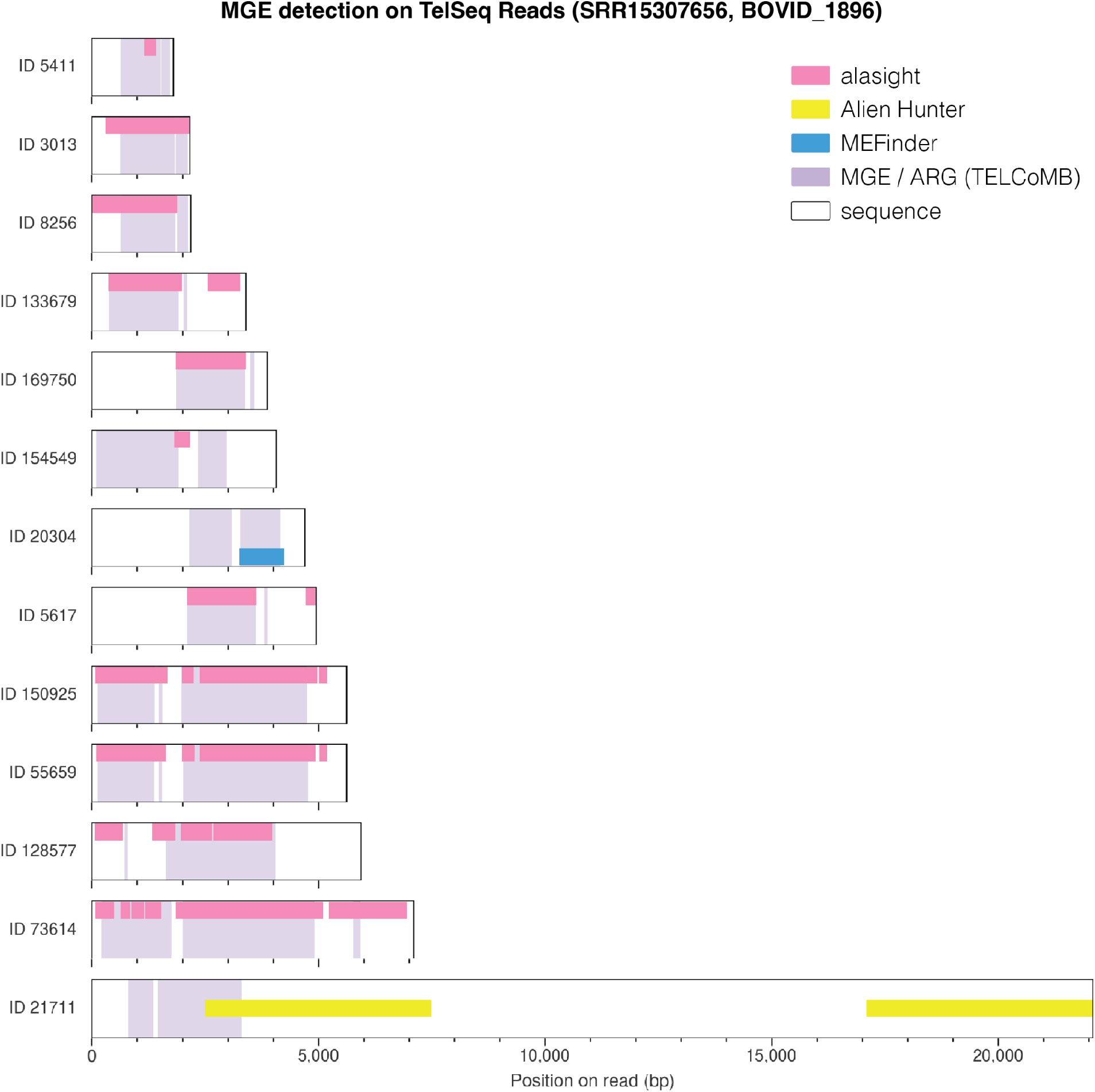
Alasight (96% ANI), MobileElementFinder (MEFinder), and Alien Hunter performance against TELCoMB ground truth (purple), visualized. Alasight hits (pink) mostly constitute a slight superset of ground truth MGEs and ARGs. geNomad flagged no regions on these reads.

## References

1. Smith, T.F. & Waterman, M.S. Identification of common molecular subsequences. Journal of molecular biology, 147(1), 195–197 (1981).

2. Altschul, S.F., Gish, W., Miller, W., Myers, E.W. & Lipman, D.J. Basic local alignment search tool. Journal of molecular biology, 215(3), 403–410 (1990).

3. Kent, W.J. BLAT—the BLAST-like alignment tool. Genome research, 12(4), 656–664 (2002).

4. Kiełbasa, S.M., Wan, R., Sato, K., Horton, P. & Frith, M.C. Adaptive seeds tame genomic sequence comparison. Genome research, 21(3), 487–493 (2011).

5. Myers, G., Durbin, R. & Zhou, C. FastGA: fast genome alignment. Bioinformatics advances, 5(1), vbaf238 (2025).

6. Li, H. Minimap2: pairwise alignment for nucleotide sequences. Bioinformatics, 34(18), 3094–3100 (2018).

7. Frost, L.S., Leplae, R., Summers, A.O. et al. Mobile genetic elements: the agents of open source evolution. Nature reviews Microbiology, 3(9), 722–732 (2005).

8. Johansson, M.H.K., Bortolaia, V., Tansirichaiya, S. et al. Detection of mobile genetic elements associated with antibiotic resistance in salmonella enterica using a newly developed web tool: MobileElementFinder. Journal of Antimicrobial Chemotherapy, 76(1), 101–109 (2021).

9. Lao, J., Lacroix, T., Guédon, G. et al. ICEscreen: a tool to detect Firmicute ICEs and IMEs, isolated or enclosed in composite structures. NAR Genomics and Bioinformatics, 4(4), lqac079 (2022).

10. Néron, B., Littner, E., Haudiquet, M., Perrin, A., Cury, J. & Rocha, E.P.C. IntegronFinder 2.0: Identification and analysis of integrons across bacteria, with a focus on antibiotic resistance in Klebsiella. Microorganisms, 10(4), 700 (2022).

11. Camargo, A.P., Roux, S., Schulz, F. et al. Identification of mobile genetic elements with geNomad. Nature Biotechnology, 42(8), 1303–1312 (2024).

12. Vernikos, G. & Parkhill, J. Interpolated variable order motifs for identification of horizontally acquired DNA: revisiting the Salmonella pathogenicity islands. Bioinformatics, 22(18), 2196–2203 (2006).

13. Zhang, X.B., Oualline, G., Shaw, J. & Yu, Y.W. skandiver: a divergence-based analysis tool for identifying intercellular mobile genetic elements. Bioinformatics, 40(Supplement_2), ii155–ii164 (2024).

14. Shaw, J. & Yu, Y.W. Fast and robust metagenomic sequence comparison through sparse chaining with skani. Nature methods, 20(10), 1661–1665 (2023).

15. Ahmed, O., Rossi, M., Kovaka, S., Schatz, M.C., Gagie, T., Boucher, C. & Langmead, B. Pan-genomic matching statistics for targeted nanopore sequencing. iScience, 24(6) (2021).

16. Li, H. Aligning sequence reads, clone sequences and assembly contigs with BWA-MEM. arXiv preprint arXiv:1303.3997 (2013).

17. Varki, R., Rossi, M., Ferro, E., Oliva, M., Garrison, E., Langmead, B. & Boucher, C. Accurate short-read alignment through r-index-based pangenome indexing. Genome research, 35(7), 1609–1620 (2025).

18. Gibson, S. & Yu, Y.W. Incorporating indel channels into average-case analysis of seed-chain-extend. Bioinformatics, 42(Supplement_1), btag312 (2026).

19. Parks, D.H., Chuvochina, M., Rinke, C. et al. GTDB: an ongoing census of bacterial and archaeal diversity through a phylogenetically consistent, rank normalized and complete genome-based taxonomy. Nucleic Acids Research, 50(D1), D785–D794 (2022).

20. Kumar, S., Suleski, M., Craig, J.M. et al. TimeTree 5: an expanded resource for species divergence times. Molecular biology and evolution, 39(8), msac174 (2022).

21. Slizovskiy, I.B., Oliva, M., Settle, J.K. et al. Target-Enriched Long-Read Sequencing (TELSeq) Contextualizes Antimicrobial Resistance Genes in Metagenomes. Microbiome, 10(1), 185 (2022).

22. Bravo, J.E., Slizovskiy, I., Bonin, N., Oliva, M., Noyes, N. & Boucher, C. The TELCoMB protocol for high-sensitivity detection of ARG-MGE colocalizations in complex microbial communities. Current protocols, 4(10), e70031 (2024).

